# Murine norovirus mutants adapted to replicate in human cells reveal a post-entry restriction

**DOI:** 10.1101/2024.01.11.575274

**Authors:** Melissa R. Budicini, Valerie J. Rodriguez-Irizarry, Robert W. Maples, Julie K. Pfeiffer

## Abstract

RNA viruses lack proofreading in their RNA polymerases and therefore exist as genetically diverse populations. By exposing these diverse viral populations to selective pressures, viruses with mutations that confer fitness advantages can be enriched. To examine factors important for viral tropism and host restriction, we passaged murine norovirus (MNV) in a human cell line, HeLa cells, to select for mutant viruses with increased fitness in non-murine cells. A major determinant of host range is expression of the MNV receptor CD300lf on mouse cells, but additional host factors may limit MNV replication in human cells. We found that viruses passaged six times in HeLa cells had enhanced replication compared with the parental virus. The passaged viruses had several mutations throughout the viral genome, which were primarily located in the viral non-structural coding regions. While viral attachment was not altered for the passaged viruses, their replication was higher than the parental virus when entry was bypassed, suggesting the mutant viruses overcame a post-entry block in human cells. Three mutations in the viral NS1 protein were sufficient for enhanced post-entry replication in human cells. We found that the human cell-adapted MNV variants had reduced fitness in mouse BV2 cells. Although the mutant viruses had increased fitness in HeLa cells, they did not have increased fitness in mice. Overall, this work suggests that MNV tropism is not only determined by the presence of the viral receptor but also post-entry factors.

**Importance:** Viruses infect specific species and cell types, which is dictated by expression of host factors required for viral entry as well as downstream replication steps. Murine norovirus (MNV) infects mouse cells, but not human cells. However, human cells expressing the murine CD300lf receptor support MNV replication, suggesting receptor expression is a major determinant of MNV tropism. To determine whether other factors influence MNV tropism, we selected for variants with enhanced replication in human cells. We identified mutations that enhance MNV replication in human cells and demonstrated that these mutations enhance infection at a post-entry replication step. Therefore, MNV infection of human cells is restricted at both entry and post-entry stages. These results shed new light on factors that influence viral tropism and host range.

## Introduction

Some viruses have very narrow host ranges and can only infect a single species while others have a much broader host range. A major determinant of host range is the presence of the viral receptor(s), as viruses can not initiate infection without cellular entry. However, some viruses are less restricted by entry because they can bind receptors on multiple species [1]. While restriction of viruses at the level of receptor binding is well understood, restriction of viruses post-entry is less well characterized. Host restriction factors can act at any stage in the replication cycle and can limit host range [1, 2]. Here, we used a forward genetic approach to select for murine norovirus (MNV) mutants with increased host range to examine factors that impact MNV tropism.

MNV is a non-enveloped single-stranded RNA virus in the *Caliciviridae* family and is a model system for human norovirus due to its robust replication in cell culture and mice [3, 4]. MNV infection has distinct outcomes depending on the viral strain. MNV1 causes acute infections that are self-resolving in immune-competent mice but can be fatal in immunocompromised STAT1-/- mice [5]. MNV3 does not cause fatal infections but can be detected in feces for weeks after infection and in some cases the virus can persist for the lifetime of the mouse [6].

CD300lf is the receptor for MNV and its expression dictates viral tropism *in vivo* and *in vitro* [7]. Both MNV1 and MNV3 can robustly infect murine immune cells that express CD300lf such as BV2 and RAW cells [8]. When CD300lf is deleted from these cells they are no longer susceptible to MNV infection. Human cells (such as HeLa) and hamster cells (such as CHO) are minimally susceptible to MNV. However, when murine CD300lf is expressed on HeLa or CHO cells they become susceptible to MNV infection. Therefore, murine CD300lf receptor expression is a major factor in determining MNV host range [7, 8]. Receptor expression is also important for tropism *in vivo*. MNV1 infection initiates in the small intestine and spreads to other sites in the intestine as well as the mesenteric lymph nodes (MLN) and spleen. Viral replication occurs within CD300lf-expressing cells in lymphoid tissue [9, 10]. Persistent infection by MNV3 initiates in the cecum and then spreads to the small intestine, colon, and the MLN. Tuft cells, which are rare intestinal epithelial cells that express CD300lf, are the site of MNV3 replication [11, 12]. Although CD300lf is crucial, there are also host factors that can impact viral replication, such as host restriction factors and immune molecules [1, 2, 13].

Forward genetic screens with viruses have been used to uncover unique facets of replication. The RNA polymerase of RNA viruses is highly error prone due to lack of proofreading, with approximately one mutation per 10,000 nucleotides [14]. Therefore, RNA viruses exist as diverse populations that are shaped by selective pressures. Past work has used sequential passaging of many RNA viruses under diverse selective pressures to gain insight into drug resistance [15, 16], antibody recognition [17, 18], tropism [19, 20], entry factors [21, 22], and host restriction factors [23, 24].

In this study we leveraged the naturally occurring diversity of MNV populations to select for mutant viruses that have enhanced replication in human HeLa cells, a non-natural host of MNV that lacks the MNV receptor CD300lf. Viruses passaged in HeLa cells had multiple mutations across the genome and included both synonymous and non-synonymous mutations. The passaged viruses had increased HeLa cell infection due to an increase in replication at a post entry step. We identified three mutations in the NS1 protein, I45F, D94E, and L231P, that may play a role in overcoming the post-entry restriction in HeLa cells. Viruses with enhanced replication in human cells had reduced fitness in murine BV2 cells indicating a fitness tradeoff for this adaptation *in vitro*. Overall, we determined that MNV can adapt to replicate in cells from a non-natural host. Additionally, our results reveal a post-entry replication restriction for MNV in human cells, suggesting factors beyond receptor availability determine MNV tropism.

## Results

### Adaptation of MNV to HeLa cells

We performed serial passages of MNV1 in HeLa cells to determine whether replication in cells from a non-natural host could increase over time. The serial passaging was done in duplicate, resulting in two similar but independent lineages. We simultaneously performed the passaging in BV2 cells, a non-selective environment, to determine if any mutations were a result of the selective HeLa cell environment or simply genetic drift from cell culture passaging. We infected BV2 and HeLa cells at MOI 0.5 with a low passage stock of MNV1 (denoted P0) and after 12 hours intracellular viral progeny were harvested by freeze-thaw cycles. We took half of the sample and amplified it using BV2 cells to generate enough virus for the next round of HeLa cell passaging (Fig. 1A). After 24 hours the BV2 cells were collected and the intracellular virus was released by freeze thawing. Half of the sample was then used to infect HeLa cells at an unknown MOI for the second round of passaging. After 6 rounds of passaging (denoted P6) and amplification we infected HeLa cells at MOI 0.5 with the passaged viruses to determine if they could replicate better than the P0 virus. After 24 hours virus was harvested and titer was determined by plaque assay using BV2 cells. We found both HeLa cell passaged stocks (P6.1 and P6.2) had increased viral yield as compared with P0 (Fig. 1B) suggesting MNV adaptation to HeLa cells.

**Figure 1.**
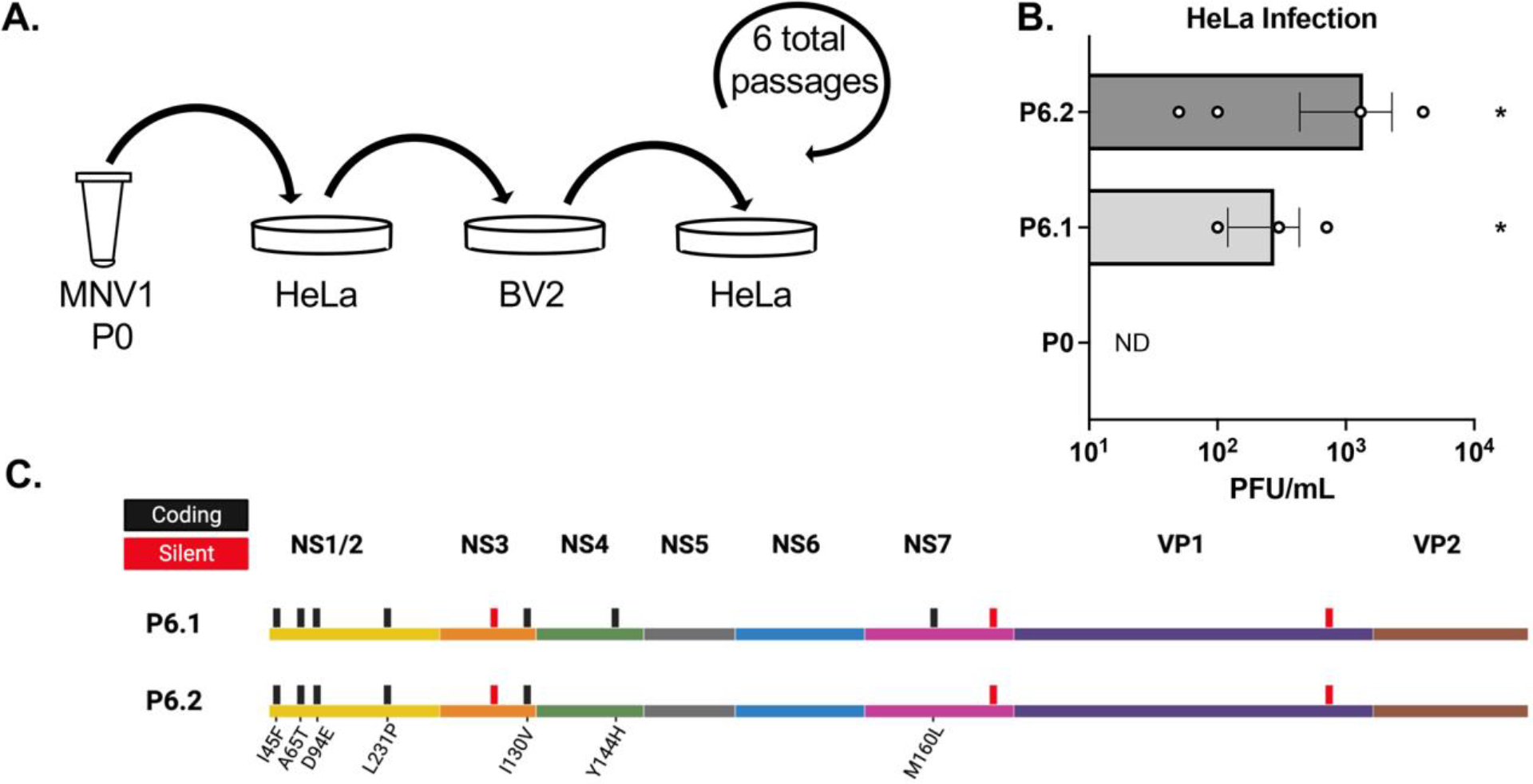
Adaptation of MNV to HeLa cells. (A) WT MNV1 virus (P0) was sequentially passaged in HeLa cells with an amplification in BV2 cells between each passage for a total of 6 passages. (B) HeLa-passaged viruses have increased replication in HeLa cells. Two independent lineages of HeLa-passaged virus (P6.1 and P6.2) and P0 were used to infect HeLa cells at MOI 0.5 for 24 hours and progeny virus was quantified by plaque assay on BV2 cells. Data are mean +/- SEM (n=4). *, *P* <0.05; Kruskal-Wallace with Dunn’s multiple comparison test with P0. ND, not detected. (C) Mutation summary from consensus sequencing of HeLa-passaged viruses. Black bars indicate coding mutations with amino acid changes underneath and red bars indicate silent mutations.

To determine if there were mutations that facilitated replication in HeLa cells we performed consensus sequencing. Viral genomes from both the BV2 and HeLa passaged viruses were amplified by RT-PCR and sequenced. We found that HeLa passaged P6.1 had 10 mutations and P6.2 had 8 mutations (Fig. 1C, Table 1). The mutations included both silent and coding mutations that spanned many regions of the viral genome. However, there was a concentration of coding mutations in the nonstructural 1/2 (NS1/2) coding region. No mutations were found in the BV2 passaged stocks after 6 passages, which suggests the mutations in the HeLa passaged stocks were a result of the specific selective pressure and not general cell culture adaptation.

**Table 1.** Mutations found in HeLa cell passaged viruses and plaque purified viruses via consensus sequencing. Gray shading denotes mutations that were cloned into new virus backbones and tested individually or in combination.* indicates A-to-G or U-to-C mutations, which are the types of mutations potentially induced by ADAR editing.

|  | Protein | Mutation | Original Codon | New Codon | AA Number | AA Change | P6 Lineage 1 | P6 Lineage 2 | P6.1.1 | P6.1.5 | 6.1.13 | P6.1.19 | 6.2.6 | 6.2.11 | 6.2.13 | 6.2.16 |
| --- | --- | --- | --- | --- | --- | --- | --- | --- | --- | --- | --- | --- | --- | --- | --- | --- |
|  | NS1/2 | Coding | ATT | TTT | 45 | I45F | X | X | X | X | X | X | X | X | X | X |
| * | NS1/2 | Coding | CTT | CCT | 60 | L60P |  |  | X |  | X | X |  | X |  |  |
|  | NS1/2 | Coding | GCG | ACG | 65 | A65T | X | X |  | X |  |  | X |  |  | X |
|  | NS1/2 | Coding | GAT | GAG | 94 | D94E | X | X | X | X | X | X | X | X | X | X |
|  | NS1/2 | Silent | TGC | TCT | 221 |  |  |  |  |  | X | X |  | X | X |  |
| * | NS1/2 | Coding | CTG | CCG | 231 | L231P | X | X | X | X | X | X | X | X | X | X |
|  | NS1/2 | Silent | CTG | TTG | 285 |  |  |  |  |  |  |  | X |  |  |  |
| * | NS1/2 | Coding | ATC | GTC | 334 | I334V |  |  |  |  |  |  | X |  |  | X |
|  | NS3 | Silent | AAC | AAT | 130 |  |  |  | X |  |  |  |  |  |  |  |
|  | NS3 | Silent | ACC | ACA | 170 |  | X | X | X | X | X | X | X |  | X | X |
| * | NS3 | Coding | ATC | GTC | 340 | I340V | X | X | X | X |  |  |  | X | X |  |
| * | NS4 | Coding | TAC | CAC | 144 | Y144H | X |  |  | X |  |  |  | X |  |  |
| * | NS4 | Coding | GAC | GGC | 146 | D146G |  |  |  |  | X | X | X |  |  | X |
| * | NS4 | Coding | GAT | GGT | 156 | D156G |  |  | X |  | X |  |  |  |  |  |
|  | NS7 | Coding | ATG | CTG | 160 | M160L | X |  |  |  |  | X |  |  |  |  |
|  | NS7 | Silent | GTG | GTT | 349 |  | X | X | X | X | X | X | X | X | X | X |
| * | VP1 | Coding | TTC | CTC | 307 | F307L |  |  |  |  |  |  | X |  |  | X |
| * | VP1 | Silent | GAA | GAG | 494 |  | X | X | X | X | X | X | X | X | X | X |

### Plaque-purified viruses from HeLa-passaged stocks have increased titer in HeLa cell infections and contain several mutations

Because there were too many mutations to individually examine, we performed further genetic analysis to identify the most relevant mutations for the HeLa cell replication phenotype. Due to our use of consensus sequencing of individual PCR products, there was a chance that not all of the mutations that we identified existed on a single viral genome and could be present at varying levels in the total population. Therefore, to determine if there was a smaller set of mutations that were sufficient for the phenotype, we plaque-purified viruses from the passaged stocks, determined their replication phenotypes in HeLa cells, and sequenced them (Fig. 2A). Because each plaque is initiated by a single founder virus, isolating plaque-purified populations allowed us to perform consensus sequencing on a viral population that is derived from a single founder virus. We diluted both P6.1 and P6.2 and plated them on BV2 cells and waited for plaques to form. After isolating well separated individual plaques (17 plaques from P6.1 and 19 plaques from P6.2), we amplified the viruses in BV2 cells. HeLa cells were infected with all of the plaque-purified viruses and we determined the titer of the viral progeny after 24 hours. We found all of the plaque-purified viruses had detectable titer whereas P0 had no detectable titer. There was a large range of titers from the different plaque-purified stocks, with P6.2.6 and P6.6.16 having the highest yield (Fig. 2B). We selected 4 plaques from each parent lineage for genetic analysis. Viral genomes from plaque-purified viruses P6.1.1, P6.1.5, P6.1.13, P6.1.19, P6.2.6, P6.2.11, P6.2.13, and P6.2.16 were amplified by RT-PCR and consensus sequences were determined. The sequencing revealed that all of the plaque-purified viruses had eight or more mutations (Table 1). Surprisingly, similar to the passaged stocks, we found very few capsid mutations which might have been expected from a selection in cells lacking the viral receptor.

**Figure 2.**
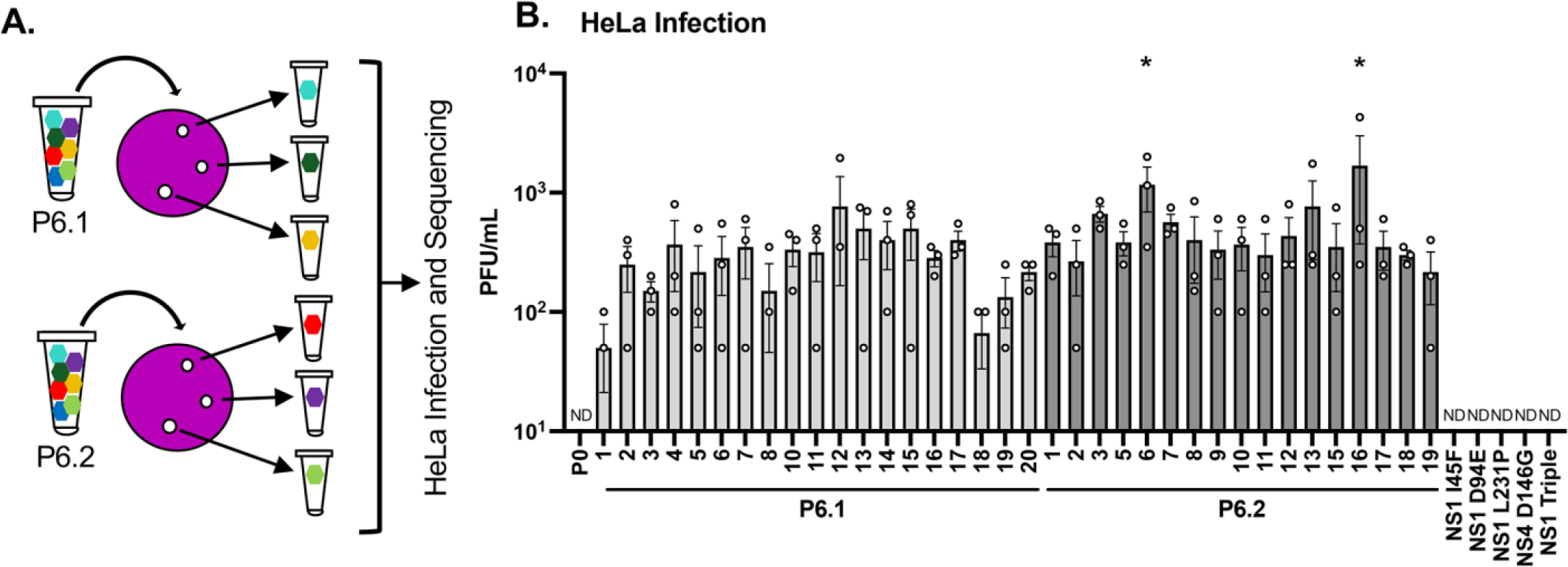
Viruses plaque purified from HeLa-passaged stocks have increased titer in HeLa cell infections. (A) Picking individual virus plaques from HeLa passaged virus. We picked 19 plaques from P6.1 and 17 plaques from P6.2 and they were amplified in BV2 cells to generate plaque purified virus stocks derived from a single founder virus. (B) Plaque-purified viruses have increased replication in HeLa cells, but viruses with individual cloned mutations do not. HeLa cells were infected with plaque-purified viruses and single or triple mutant cloned viruses (MOI 0.5). Viral titer was determined after 24 hours by plaque assay in BV2 cells. Data are mean+/- SEM (n=3). *, *P* <0.05; one-way ANOVA with Fisher’s multiple comparison test with PO. ND, not detected.

### Individual mutations are not sufficient for HeLa adaptation

Given the relatively large number of mutations present in HeLa-adapted viruses, we decided to narrow our focus primarily to mutations in NS1 since several were common among plaque-purified viruses with enhanced replication. MNV NS1 is a multifunctional protein implicated in replication and tropism *in vitro* and *in vivo*. NS1/2 is cleaved from the larger ORF1 viral polyprotein by the viral protease and then host caspase 3 cleaves NS1/2 to generate NS1 and NS2 [25]. NS1/2 interacts with host VAMP-associated protein A (VAPA), which stimulates a replication step after viral entry but before RNA synthesis [26] and NS1/2 may drive replication complex formation by recruiting cellular membranes [27]. NS1 can be secreted from cells and plays a role in interferon resistance *in vivo* [28]. Furthermore, NS1 is a major determinant of cellular tropism and persistence in mice. An acute strain of MNV that normally replicates in immune cells can be converted into a persistent strain that replicates in epithelial (Tuft) cells by expressing NS1 of the persistent strain [29].

Importantly, the presence of a glutamate instead of aspartate at amino acid 94 of NS1 is sufficient for viral persistence and replication in Tuft cells [30]. We decided to focus on three mutations in NS1 that were found in the passaged lineages and all eight of the plaque-purified viruses (NS1 I45F, D94E, and L231P) as well as one mutation D146G in NS4 that was present in many of the plaque-purified viruses with high viral yield (Table 1, gray highlighted rows). Note that the D94E mutation influences viral tropism (immune cells vs. epithelial cells) and persistence in mice [30]; however, its effect on replication in human cells was unknown.

To determine if any of the individual mutations that were common in all of the plaque-purified HeLa passaged viruses were sufficient for HeLa cell infection, we cloned them into a MNV1 infectious clone using site-directed mutagenesis and generated virus stocks. We made individual single mutations for NS1 I45F, NS1 D94E, NS1 L231P, and NS4 D146G. We also made a triple mutant with all three of the NS1 mutations (NS1 triple). We then infected HeLa cells at MOI 0.5 with all of the cloned mutants and P0 and after 24 hours quantified the viral progeny by plaque assay. Although these mutants were viable in BV2 cells, there was no detectable titer in HeLa cells with any of the single mutants or the triple mutant (Fig. 2B). From this we concluded that a larger set of mutations is likely needed for HeLa cell adaptation, so we continued our further analyses with the eight selected plaque-purification viruses as well as the cloned mutants.

### HeLa-passaged viruses do not have increased attachment to HeLa cells

To determine which step of the viral replication cycle was altered by HeLa cell adaptation we started by examining cell attachment, the first step of the viral replication cycle. We performed attachment assays with P0, plaque-purified viruses, and cloned mutant viruses. Each viral strain was incubated with HeLa cells at an MOI 2 for one hour at 4°C which facilitates viral binding but not entry, followed by washing and quantification of cell-associated virion RNA using qRT-PCR. P0 and all of the viruses tested had similar low levels of attachment to HeLa cells comparable to the no cell negative control, indicating that all of the viruses had minimal binding to HeLa cells (Fig. 3). To confirm this experiment is sufficient to detect MNV binding we also incubated P0 virus with BV2 cells and CD300lf HeLa cells that express the MNV receptor. The P0 virus had significantly higher levels of attachment to these cells (Fig. 3). These results suggest the HeLa cell adaptation did not improve the virus’ ability to attach to HeLa cells and suggests that the viruses may be entering through a CD300lf-independent mechanism.

**Figure 3.**
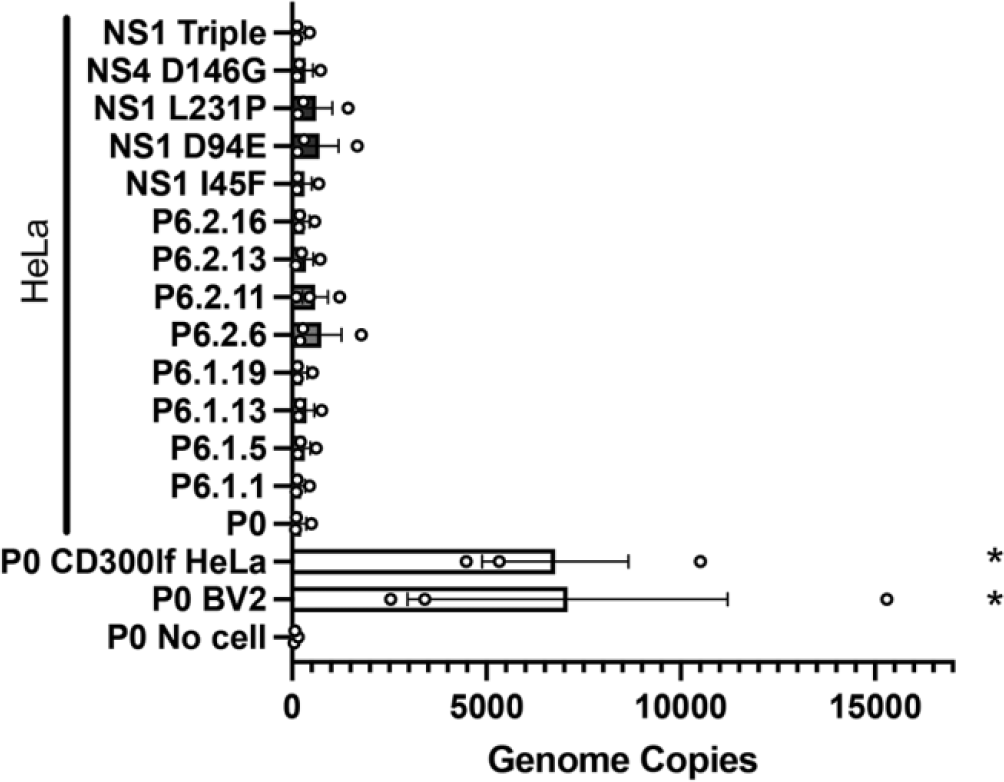
HeLa-passaged viruses do not have increased attachment to HeLa cells. BV2, CD300lf-HeLa, or HeLa cells were incubated with P0, plaque purified, or cloned mutant virus (MOI 2) for 1 hour at 4°C. Cells were washed and cell-associated vRNA was isolated and quantified by qRT-PCR. Data are mean+/- SEM (n=3). *, P <0.05; one-way ANOVA with Fisher’s multiple comparison test with P0.

### HeLa-adapted viruses overcome a post-entry restriction

Since there was no attachment advantage amongst the HeLa-passaged viruses, we hypothesized that they have a replication advantage post-entry. To test this, we performed an entry bypass experiment by transfecting HeLa cells with viral RNA (vRNA) and quantifying viral yield by plaque assay. Because MNV is a positive-sense single stranded RNA virus, transfecting RNA is sufficient to initiate viral translation, replication, and production of infectious virus in permissive cells. Therefore, we used this method to determine if the HeLa-adapted viruses have replication advantages in HeLa cells post-entry. vRNA from P0, plaque-purified viruses, and cloned mutant viruses was isolated, quantified by qRT-PCR, and equivalent amounts of RNA were transfected into HeLa cells. After 12 hours, the viral progeny from both the cells and supernatant were quantified by plaque assay. We found that most of the plaque-purified viruses had an increase in viral yield as compared with P0. Additionally, the NS1 triple mutant had similar yield to the plaque-purified viruses (Fig. 4A). We also confirmed the post-entry replication advantage in HeLa cells by infecting CD300lf expressing HeLa cells. Because these cells express the MNV receptor they do not have entry restriction. We infected CD300lf HeLa cells with P0 virus, plaque-purified viruses, and cloned mutant viruses at an MOI of 0.5 and quantified the viral progeny after 24 hours by plaque assay. The trends in these data were similar to the entry-bypass experiment with all of the plaque-purified viruses having a ∼100 fold increase in viral yield over P0. The NS1 D94E, L231P, and triple mutant had similar yield to the plaque-purified viruses (Fig. 4B). Taken together these data suggest HeLa cells restrict MNV replication post-entry and our HeLa adapted viruses are able to overcome this restriction.

**Figure 4.**
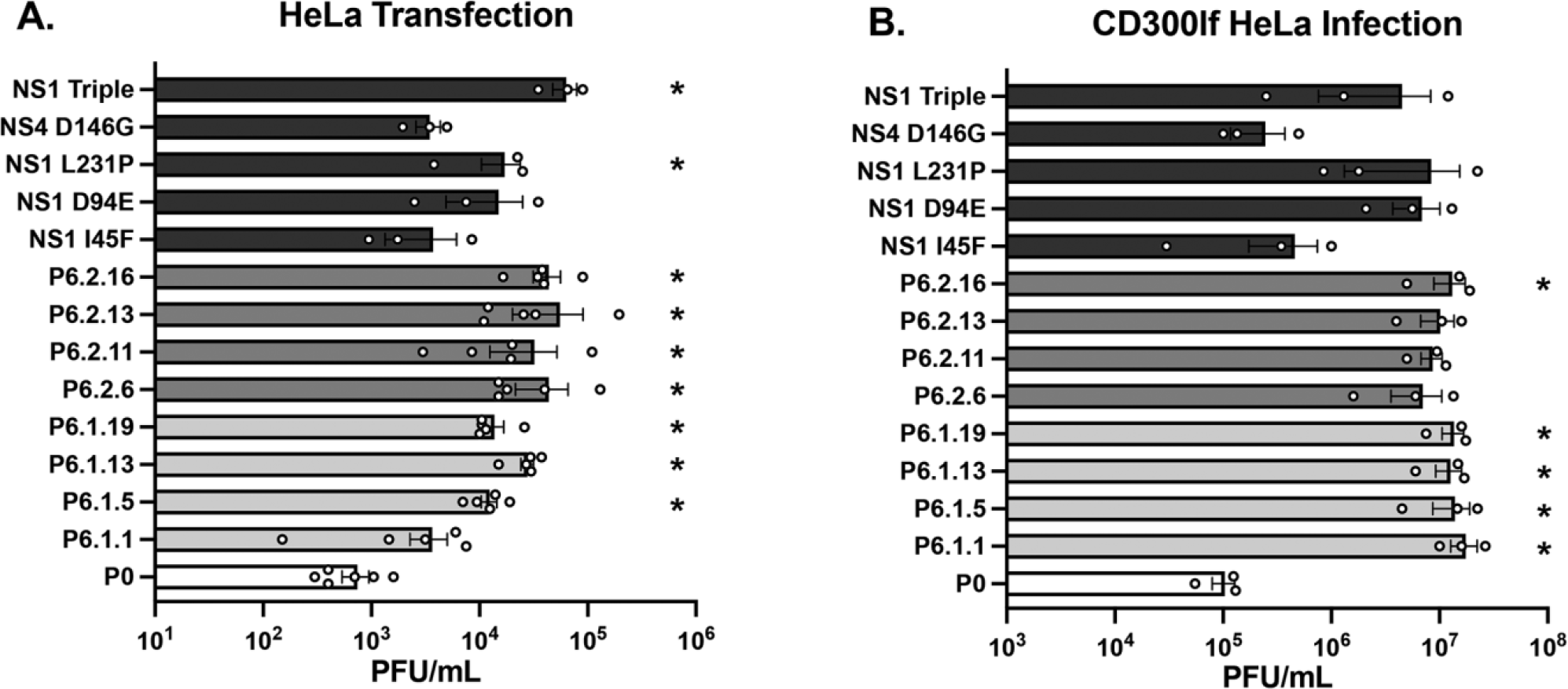
HeLa-adapted viruses overcome post-entry restriction. (A) HeLa cells were transfected with viral RNA from P0, plaque purified, and cloned mutant viruses. After 12 hours cells and supernatant were collected. Infectious virus was quantified by plaque assay in BV2 cells. Data are shown as mean+/- SEM (n=3-6). *, P <0.05; Kruskal-Wallace with Dunn’s multiple comparison test with P0. (B) CD300If-HeLa cells were infected with either P0, plaque purified, or cloned mutant virus (MOI 0.5). Viral titer was determined after 24 hours using BV2 cells. Data are shown as mean+/- SEM (n=3). *, P <0.05; one-way ANOVA with Fisher’s multiple comparison test with P0.

### HeLa-passaged virus and NS1 triple mutant have increased viral protein production in HeLa cells

We next determined whether the post-entry MNV restriction in HeLa cells limits viral protein production and whether the HeLa-adapted viruses had higher protein yields. We infected BV2 cells with P0 and 6.2.16 virus and CD300lf HeLa cells with P0, P6.2.16, and NS1 triple mutant viruses at an MOI of 0.5. After 24 hours cell lysates were generated and viral protein levels were detected by western blot with an anti-NS1 antibody. In BV2 cells we detected viral protein for both P0 and P6.2.16 virus as expected (Fig. 5). However, in the CD300lf HeLa cells there was no NS1 band detected for P0 while an NS1 band was detected for P6.2.16 and NS1 triple mutant (Fig. 5). Since MNV protein levels reflect both initial translation from incoming RNA as well as translation of nascent RNAs following RNA synthesis, the lack of detectable NS1 for P0 virus could reflect defects in translation, RNA replication, or both. Regardless, these results indicate that restriction of MNV in HeLa cells limits viral protein production and HeLa-adapted viruses overcome this restriction.

**Figure 5.**
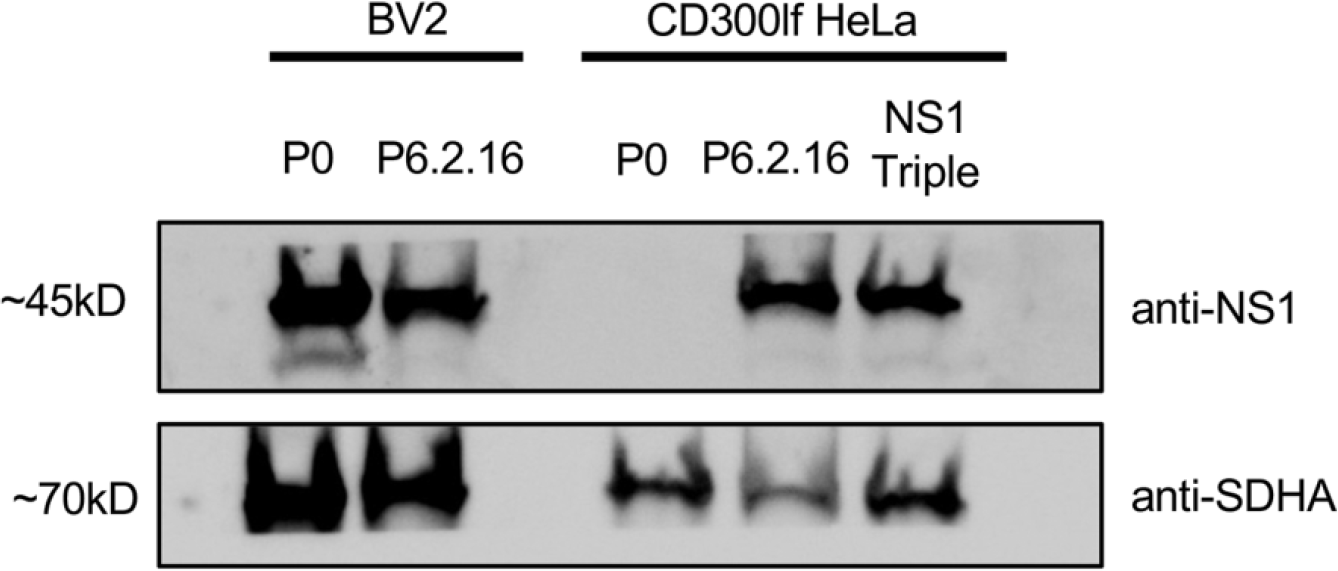
HeLa-passaged virus and NS1 triple mutant virus have increased viral protein production in HeLa cells. Representative western blot from BV2 and CD300lf HeLa cells infected with P0, 6.2.16, or NS1 triple mutant (MOI 0.5). At 24 hours post-infection cells were lysed and blots were probed with anti-NS1 antibody to detect viral protein or anti-SDHA antibody as a loading control.

### HeLa-passaged viruses have decreased fitness in BV2 cells

Because the HeLa-passaged viruses accumulated several mutations during HeLa cell adaptation, we wondered whether there was a fitness cost in a different environment. To test this, we infected BV2 cells with P0, plaque-purified, and cloned mutant viruses at MOI 0.5. After 24 hours we determined viral yield by plaque assay. The plaque-purified viruses replicated to significantly lower levels than the P0 virus, indicating that the HeLa-adapted viruses have reduced fitness in BV2 cells. Interestingly, the NS1 triple mutant virus, which is sufficient to overcome the post-entry replication restriction in HeLa cells, does not have reduced fitness in BV2 cells (Fig. 6A). This may indicate only specific mutations or a larger number of mutations are responsible for the reduced fitness. We also assessed the replication capacity of the adapted viruses in BV2 cells by calculating the vRNA to PFU ratio after infecting BV2 cells. To do this we infected BV2 cells, collected samples at 24 hpi, and used half of each sample for plaque assays using BV2 cells and used the other half of each sample for RNA isolation and quantification of viral genomes by qRT-PCR. We then divided the number of genomes per sample by the titer. A higher RNA:PFU ratio indicates the virus has reduced efficiency of replication. We found that two of the plaque-purified viruses, P6.2.6 and P6.2.16, had significantly higher RNA:PFU ratios than P0 indicating a fitness cost in BV2 cells. Although not statistically significant, many of the other plaque-purified viruses also had higher ratios than P0 while most of the cloned mutants did not (Fig. 6B). Overall this indicates the mutations that lead to HeLa cell adaptation have fitness costs in other cell environments.

**Figure 6.**
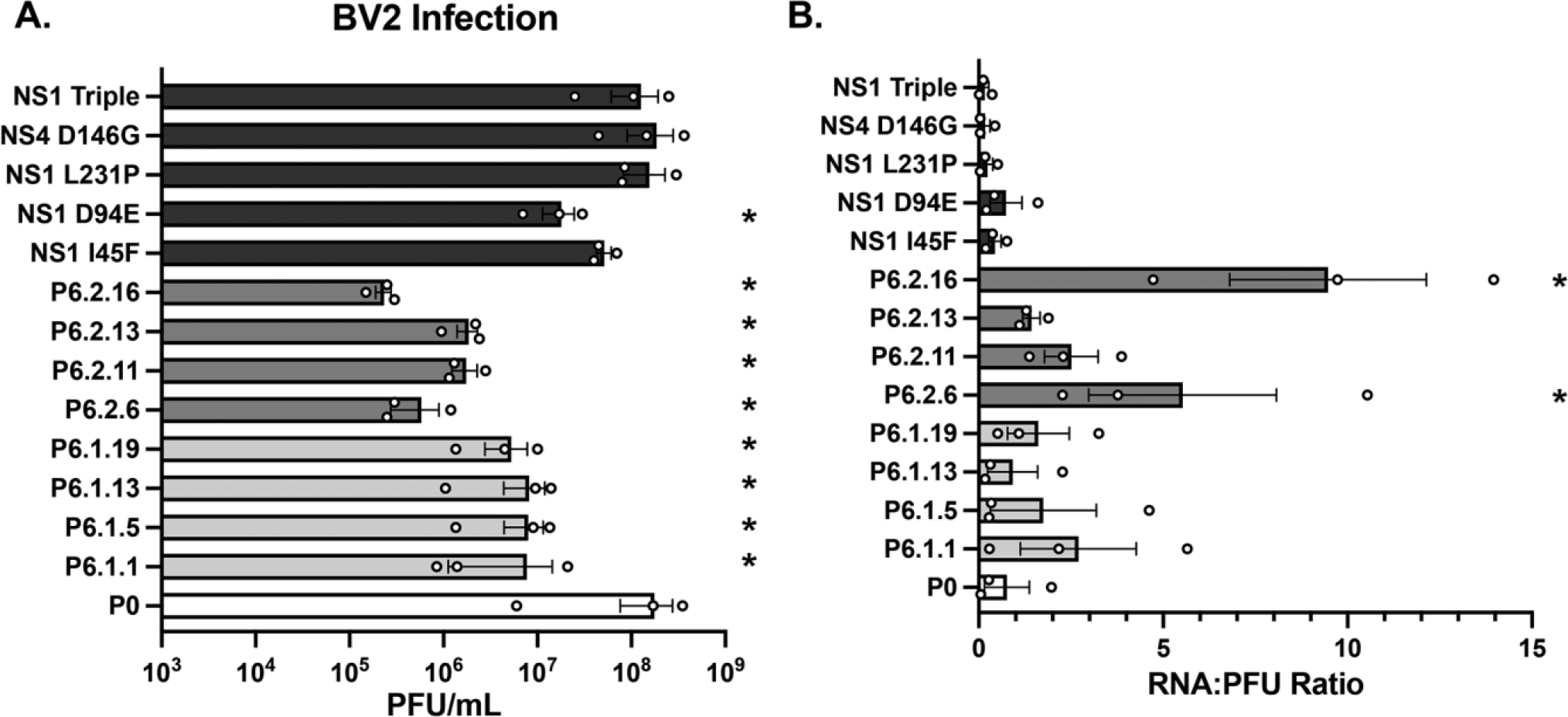
HeLa-passaged viruses have decreased fitness in BV2 cells. (A) Passaged viruses have decreased replication in BV2 cells. BV2 cells were infected with P0, plaque purified, or cloned mutant virus (MOI 0.5). Viral titer was determined after 24 hours using BV2 cells. Data are shown as mean+/- SEM (n=3). *, P <0.05; one-way ANOVA with Fisher’s multiple comparison test with P0. (B) RNA to PFU ratio is increased in some passaged viruses. Half of the infected BV2 cells were collected in Trizol and viral RNA was quantified using qRT-PCR, and the other half was used for plaque assays in BV2 cells. The RNA:PFU ratio represents the viral genome number divided by the viral titer for each sample. Data are shown as mean+/- SEM (n=3). *, P <0.05; one-way ANOVA with Fisher’s multiple comparison test with P0.

### HeLa-adapted viruses do not have altered shedding in mice

Since the HeLa-passaged viruses have reduced fitness in BV2 cells but enhanced fitness in HeLa cells, we wondered whether HeLa adaptation confers a fitness cost *in vivo*. To test this we used 1 x 10^6^ PFU virus to perorally infect male and female C57BL/6 mice. The mice were infected with six viral strains, MNV1 (P0, an acute strain), MNV3 (a persistent strain), CR6 (a persistent strain), P6.2.16, NS1 D94E, and NS1 triple mutant virus. Feces were collected on day 0 before infection to confirm that the mice were not infected with MNV prior to the experiment. Fecal pellets were also collected on day 3, 7, 14, and 21 after infection to monitor replication and shedding. Viral genomes were quantified using qRT-PCR. As expected, we found that MNV3 and CR6 had increased titer and prolonged shedding since they are persistent strains of MNV. We found there were no significant differences between the parental MNV1 strain and the HeLa-adapted viruses (P6.2.16, NS1 D94E, and NS1 triple) indicating the HeLa adaptation mutations do not alter shedding in WT mice compared to their parental virus (Fig. 7). Previously, Nice et al. demonstrated that a NS1 D94E mutation in an acute MNV strain induced viral persistence [31]. Curiously, although this same NS1 D94E mutation is present in all three of our adapted viruses, it did not cause persistent, prolonged shedding. Overall, our results indicate that the mutations resulting from HeLa cell adaptation do not increase infection in mice.

**Figure 7.**
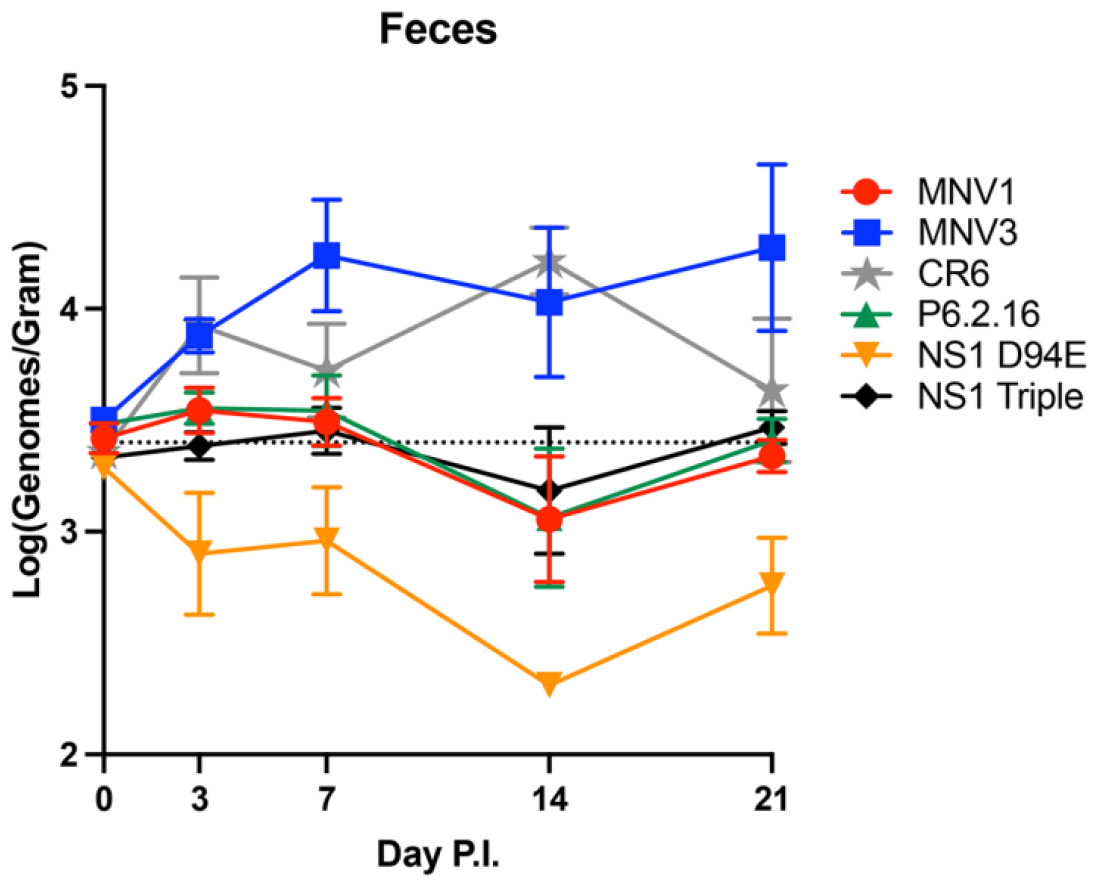
HeLa-passaged virus or cloned mutants do not have altered shedding in WT mice. Both male and female mice were orally infected with 1 x 10^6^ PFU of MNV1 (P0/acute strain), MNV3 (persistent strain), CR6 (persistent strain), P6.2.16, NS1 D94E, and NS1 Triple mutant viruses. Feces was collected at day 0 before infection and was also collected at days 3, 7, 14, and 21 post infection. MNV genomes were quantified by qRT-PCR. Data are mean +/- SEM (n=6 over 2 independent experiments). Dashed line indicates the detection threshold.

## Discussion

In this work we examined factors that impact MNV tropism by selecting for and characterizing variants with increased replication in human cells. It is well known that the viral receptor, CD300lf, is important for tropism both *in vivo* and *in vitro*. However, other factors that determine tropism are less understood. In this study we found that HeLa-adapted viruses did not have increased attachment to HeLa cells. Instead, we identified mutations that allowed these viruses to overcome a post entry restriction present in HeLa cells, with increased viral protein synthesis. Overall, we found that viral tropism is not determined exclusively by viral receptor expression but also by cell-dependent restriction post-entry.

Previously our lab and others have used viral passaging in restrictive environments to select for RNA viruses with various traits such as faster replication, drug resistance, increased thermostability, and altered plaque size [32-35]. Most of these viral passaging studies have resulted in a small number of mutations and often the phenotype of interest can be traced back to a single mutation. Therefore, it was interesting that after passaging MNV in HeLa cells we uncovered eight to ten mutations, and we could not entirely attribute the increased replication phenotype to one specific mutation.

It is unclear what led to the large number of mutations in our viruses under these passaging conditions. One hypothesis is that the mutation frequency alone was insufficient to overcome both entry and post-entry restriction in HeLa cells and the viral genome may have been subjected to host RNA editing mechanisms to introduce additional mutations. One mechanism that may have introduced additional mutations is the host ADAR protein, a cellular factor that edits RNA as part of the innate immune response and post transcriptional modification [36-41]. There is evidence that other viral genomes such as SARS-CoV-2 may have undergone editing by host proteins [42]. One genetic signature indicative of ADAR mutations is A-to-G or U-to-C mutations, and these types of changes represent half of the mutations that we uncovered in the plaque-purified mutants [43] (Table 1, mutations with *). Further analysis would be required to determine if this host editing mechanism may have played a role in mutating the virus during HeLa cell passaging.

Knowing that HeLa cells do not express the MNV receptor CD300lf, we expected that the main restriction for MNV in HeLa cells would be at the level of entry. We hypothesized that lack of the viral receptor would result in capsid mutations that may alter viral attachment and entry. It was surprising when we found most of the mutations in the passaged viruses were located in the non-structural proteins. This led us to the discovery that the passaged mutants increase viral replication when entry is bypassed or when CD300lf is expressed in HeLa cells (Figs. 4 and 5). We also found that none of the mutations in the non-structural proteins altered cell attachment. This suggests the virus must have been able to (perhaps inefficiently) enter cells during the passaging. However, there are no known CD300lf-independent entry mechanisms for MNV in cultured cells. Interestingly, Kennedy et al. recently demonstrated passive uptake of MNV in the ileum of neonatal mice and replication-independent shedding, suggesting passive absorption through a largely CD300lf-independent mechanism [44]. It was interesting that although the mutations that we cloned individually into the MNV backbone increased viral replication when entry was bypassed (Fig. 1A) they had no impact on HeLa infection even though the plaque-purified viruses did increase infection (Fig. 2B). This may indicate that some of the other mutations found in the plaque-purified viruses that we did not clone individually may play a role in viral entry. Future investigation is required to determine the mechanism of viral entry into HeLa cells that do not express CD300lf.

Although we did not find a single mutation with the same level of replication as the plaque-purified viruses when entry was bypassed, we did find that the NS1 triple mutant could replicate to the same level. Therefore, NS1 may be involved in the post-entry restriction of MNV in human cells. We also found the NS1 triple mutant had increased viral production in CD300lf HeLa cells as compared with P0. Previous studies showed NS1 plays a role in replication complex formation and that NS1 is a determinant of cell tropism *in vivo* [26, 27, 29]. These previous reports and our data suggest a potential function for NS1 in determining cell tropism specifically at the step of viral replication.

In previous selections with other RNA viruses, adaptation to one environment can have fitness trade-offs that reduce fitness in other environments [32, 33, 45, 46]. In our study we found that all the plaque-purified viruses from the HeLa-passaged lineages had decreased replication in BV2 cells. Interestingly 4/5 of the cloned mutations did not reduce fitness in BV2 cells (Fig. 6A). This may indicate that the number of mutations may have an impact on fitness in BV2 cells. Although the passaged viruses had increased fitness in HeLa cells, they did not have increased fitness in mice. Overall, these results indicated that multiple mutations contribute to fitness and host range and that forward genetic approaches can reveal new insights into viral tropism.

## Materials and Methods

### Cells and Viral Stocks

BV2 cells were maintained in Dulbecco’s Modified Eagle Medium (DMEM) with 10% fetal bovine serum, 1% HEPES, and 1% penicillin-streptomycin. HeLa cells were maintained in DMEM with 10% calf serum and 1% penicillin-streptomycin.

HeLa cells stably expressing the MNV receptor CD300lf were generated by lentivirus transduction as previously reported, in collaboration with Robert Orchard’s lab [24]. Briefly, lentivirus was generated by transfecting lentiviral vectors with packaging and expression plasmids into 293T cells. Supernatants were filtered and added to the same HeLa cells used for the passaging. The cells expressing CD300lf were selected with puromyocin. Individual colonies were picked and expanded, and each was tested for functional CD300lf receptor expression by quantifying relative yield after infection with MNV1. The clone with the highest viral yield was then used as our CD300lf HeLa cell line.

MNV-1.CW3 (MNV1) [47] was generated by transfecting HEK293T cells with an infectious clone plasmid followed by two rounds of amplification in BV2 cells to generate high titer viral stocks. Viral stocks were stored at -80°C. WT MNV1 is referenced as P0 throughout.

To quantify virus, plaque assays were performed as previously described. Briefly, virus was diluted in phosphate-buffered saline supplemented with 100 μg/ml CaCl_2_ and 100 μg/ml MgCl_2_ (PBS+) and added to BV2 cells for 20-30 min at 37°C to facilitate attachment. Overlays containing 1.5% methylcellulose, MEM, 10% fetal bovine serum, 1% penicillin-streptomycin, and 1% HEPES were used and removed after 48-72 h.

### Viral Passaging Experiments

BV2 and HeLa cells were plated on 6-well plates and cells were infected with MNV1 (P0) using an MOI of 0.5. Inoculum virus was absorbed for 1 hour at 37 °C, the supernatant was aspirated, the cells were washed with PBS+, and 1 mL of fresh media was added. The infection was allowed to continue for 12 hours and then the cells were scraped into the media and the entire 1 mL sample was freeze-thawed 3 times to release the intracellular virus. Then, 500 μL of the sample was used to infect BV2 cells seeded in 6-well plates to amplify virus. The infection conditions for this amplification were the same as above except the infection was allowed to continue for 24 hours. Following amplification, 500 μL virus was then used to begin a new infection of BV2 and HeLa cells at an unknown MOI. This cycle of infection and amplification was continued for 6 passages. The selection was done in duplicate resulting in two independent lineages (P6.1 and P6.2). The P6 stocks were plaque purified to isolate individual viral populations derived from a single founder virus. For P6.1, 19 plaque-purified viral stocks were generated and for P6.2 17 stocks were generated. The plaque-purified viral stocks were amplified in BV2 cells.

### Sequencing

The virus collected from P6 and the plaque-purified viruses were used to infect BV2 cells and viral RNA was isolated with Trizol. cDNA was generated by RT-PCR with SuperScript IV reverse transcriptase (ThermoFisher) and primer 5’ TGGCGTCAGACTTTGAGAAGC 3’. PCR was performed with primer pairs that spanned the viral genome i.e. sense primer (1) 5’ GTGAAATGAGGATGGCAACG 3’ antisense (1) 5’ GGGTCCA AAAGATGTCAAA GA 3’, sense (2) 5’ GCTCAACATTCTCAACATC G 3’ antisense (2) 5’ GACCGCCTCCAGGTTGAC 3’, sense (3) GACAGGATTGAGAACAAGGG 3’ antisense (3) 5’ GTAATCATCAATGGAGTACTTG 3’, sense (4) 5’ CATGGATACACCTACCGTGA antisense (4) 5’ CTGGAACTCCAGAGCCTCAA 3’, sense (5) 5’ CACCTGGGTTGTGATTGGG 3’ antisense (5) 5’ ATGTGGTACCTGAAATTGGC 3’, sense (6) 5’ GCCAACAACATGTATGAGATG 3’ antisense (6) 5’ AGTCCTGTAATACTTTTCACCA 3’, sense (7) 5’ ACGCTGCAGGGCATCTCC 3’ antisense (7) 5’ ACGCTGCAGGGCATCTCC 3’, sense (8) 5’ T GGTCTCTGGCCGCCTTC 3’ antisense (8) 5’ CTTCCCACAGAGGCCAATTG 3’, sense (9) 5’ ACACCGCTGACGCCGCAG 3’ antisense (9) 5’ TTTTAAAATGCATCTAAATACTACT 3’. PCR products were sequenced by Genewiz from Azenta Life Sciences.

### Cloning MNV Mutants

The sequencing revealed that there were many mutations present in the both P6 lineages as well as the plaque-purified viral stocks. We decided to focus on three mutations that were present in all of the sequenced stocks and one that was present in many of the plaque-purified stocks. The mutations were cloned by Genewiz/Azenta Life Sciences using site-directed mutagenesis into a MNV1 plasmid. We then transfected the plasmids into HEK293T cells and generated viral stocks as described above. The presence of the mutations was sequence confirmed both in the plasmid (by whole plasmid sequencing by Genewiz/Azenta Life Sciences) and in the resulting viral stocks.

### Viral Infection Assay

12-well plates were seeded with 3 x 10^5^ HeLa, CD300lf-HeLa, or BV2 cells. Each cell line was inoculated with either P0, selected plaque purification stocks, or the cloned mutations at MOI 0.5 for 1 hour at 37°C. The inoculum was removed and the cells were washed with PBS+, 1 mL of fresh media was added, and cells were incubated for 24 hours at 37°C. For the HeLa and CD300lf HeLa infections the cells were pelleted and resuspended in 200 μL PBS+. To release the virus the cell pellets were freeze and thawed three times before titer was determined by plaque assay on BV2 cells. For the BV2 infection the cells were pelleted and half were resuspended in 100 μL PBS+ and used for plaque assays on BV2 cells. The remaining cells were resuspended in Trizol and viral RNA was isolated and quantified using qRT-PCR as described below. The RNA:PFU ratio was calculated by dividing the genome copy number by the titer for each sample.

### Viral Attachment Assay

6 well plates were seeded with 3 x 10^5^ BV2, HeLa, CD300lf-HeLa, and a no cell control well. The BV2, HeLa, and no cell control wells were inoculated with P0 virus at MOI 2 in 1mL cold DMEM media containing 10% FBS. HeLa cells were also infected with plaque-purified viruses and cloned mutant viruses. The virus was allowed to bind on ice at 4°C for 1 hour. The cells were washed three times with PBS+ before they were collected in 750 μL Trizol. To determine the amount of virus that bound the cells viral RNA was isolated with RxnDirect-zol RNA Miniprep Plus kit (Zymo Research) according to the manufacturers’ protocol. cDNA was generated and TaqMan quantitative PCR (qPCR) for MNV was performed in triplicate for each sample as described previously to quantify viral genomes [48]. The forward primer 5′-GTGCGCAACACAGAGAAACG-3′, reverse primer 5′-CGGGCTGAGCTTCCTGC-3′, and probe 5′-6FAM-CTAGTGTCTCCTTTGGAGCACCTA-BHQ1-3′ were used for all samples. These sequences are in the viral NS1/2 coding region, but do not overlap with any of the mutations present in our passaged viruses. MNV1 plasmid that had been linearized with FseI was used as a standard.

### Entry Bypass Assay

Viral RNA stocks were generated by infecting BV2 cells with MNV and isolating RNA with Trizol (Invitrogen) according to the manufacturers protocol. The viral RNA concentration in each stock was determined by qRT-PCR as described in binding assay to ensure that the same amount of genome copies were transfected per sample. 24 well plates were seeded with 1 x 10^5^ HeLa cells. Viral RNA (1.2 x 10^7^ genome equivalents) from P0, plaque-purified stocks, or cloned mutation stocks was transfected using Lipofectamine 2000 (Invitrogen) according to the manufacturers’ protocol. After 12 hours the cells and supernatant were collected, and the samples were freeze thawed 3 times before determining viral titer by plaque assay.

### Western Blot Detection of MNV

Either BV2 or HeLa cells in 12 well plates were inoculated with P0, P6.2.16, or NS1 triple mutant at MOI of 0.5 as described above. However, after the cells were collected at 24 hpi, the pellet was resuspended in 200 μL 4x Laemmli Sample Buffer with 10% 2-mercaptoethanol (BioRad). The samples were denatured by boiling for 20 minutes. Lysates were run on SDS-PAGE gels and transferred to PVDF membranes. The antibody used to detect MNV was a mouse monoclonal anti-NS1 (a kind gift from Robert Orchard, 1:1000). SDHA was detected as a loading control with a mouse monoclonal anti-SDHA antibody (Abcam, 1:1000).

### Mouse Experiments

All animals were handled according to the Guide for the Care of Laboratory Animals of the NIH. All mouse studies were performed at UT Southwestern (Animal Welfare Assurance no. a3472-01) by using protocols approved by the local Institutional Animal Care and Use Committee in a manner designed to minimize pain, and any animals that exhibited severe disease were euthanized immediately with isoflurane. Five-week old WT C57BL/6 mice were obtained from Jackson Laboratories (Jax#000664) immediately prior to the experiment (to limit potential exposure to any MNV in local animal facilities) and were singly housed during the experiments. Both male and female mice were used. Mice were perorally infected with 1 x 10^6^ PFU of virus. Feces were collected prior to infection (day 0) and on days 3, 7, 14, and 21 post infection. At day 21 postinfection the mice were euthanized and tissues were collected. The feces were homogenized by bead beating and viral RNA was extracted using the RNeasy Mini QIAcube kit (Qiagen). The tissues were homogenized by bead beating and viral RNA was extracted using the RxnDirect-zol RNA Miniprep Plus kit (Zymo). For all samples the viral RNA was quantified by qRT-PCR as described above.

### Ethics/Biosafety Statement

These gain-of-function selection experiments using a non-respiratory BSL2 virus were approved by the University of Texas Southwestern Institutional Biosafety Committee (IBC) prior to work. Additionally, the IBC was updated with progress throughout the project, including identification of specific mutations and approval prior to starting animal experiments.

### Statistical Analysis

All error bars represent the standard error of the mean (SEM). Specific statistical test used for each experiment is indicated in the figure legends. For all pairwise comparisons a p-value <0.05 was considered significant and represented with a single asterisk. GraphPad Prism was used to perform all statistical analyses.

## Acknowledgements

We thank Andrea Erickson for critical review of the manuscript. We are extremely grateful to Robert Orchard for sharing critical MNV reagents, protocols, and advice.

Work in J.K.P.’s lab is funded through R37 AI074668, and this work was partially funded by awards from the HHMI Faculty Scholars Program and the Burroughs Wellcome Fund Investigators in the Pathogenesis of Infectious Diseases program. M.R.B. and V.R.I were supported in part by the NIH Molecular Microbiology Training Grant T32 AI007520. R.W.M was supported in part by T32 AI005284.

